# Optimization of highly inclined Illumination for diffraction-limited and super-resolution microscopy

**DOI:** 10.1101/2023.02.17.528478

**Authors:** L. Gardini, T. Vignolini, V. Curcio, F.S. Pavone, M. Capitanio

## Abstract

In HILO microscopy, a highly inclined and laminated light sheet is used to illuminate the sample, thus drastically reducing background fluorescence in wide-field microscopy, but maintaining the simplicity of the use of a single objective for both illumination and detection. Although the technique has become widely popular, particularly in single-molecule and super-resolution microscopy, a limited understanding of how to finely shape the illumination beam and how this impact on the image quality complicates setting HILO to fit the experimental needs. In this work, we build up a simple and comprehensive guide to optimize the beam shape and alignment in HILO and to predict its performance in conventional fluorescence and super-resolution microscopy. We model the beam propagation through Gaussian optics and validate the model through far- and near-field experiments, thus characterizing the main geometrical features of the beam. Further, we fully quantify the impact of a progressive reduction of the inclined beam thickness on the image quality of both diffraction-limited and super-resolution images and we show that the most relevant impact is obtained by reducing the beam thickness to sub-cellular dimensions (< 3 μm). Based on this, we present a simple optical solution to reduce the inclined beam thickness down to 2.6 μm while keeping a field-of-view dimension suited for cell imaging and allowing an increase in the number of localizations in super-resolution imaging of up to 2.6 folds.

## INTRODUCTION

Single Molecule Localization (SML) imaging relies on the detection of a large number of photons to perform effective fitting of the intensity profile of a single light emitter and achieve nanometer localization accuracy and image resolution [1][2]. Application of SML microscopy to cell cultures and thick samples in a wide-field configuration shows a significant limitation caused by the high background from off-focus fluorescence. Total Internal Reflection (TIR) excitation is widely applied to overcome this drawback, giving access to fluorophores within about 200 nm from the glass surface [3]. By exciting such a thin portion of the sample, the background fluorescence is minimized and SML with nanometer precision is made possible on membrane proteins, lipids and other components of the plasma membrane. However, the very limited penetration depth of the evanescent wave does not allow the study of structures and dynamics occurring inside a cell or a tissue.

On the contrary, confocal microscopy allows for low background imaging inside thick samples through the confinement of the excitation beam to the volume of a focused laser beam and the rejection of off-focus fluorescence by pinhole filtering. However, despite the development of several improved designs, this technique is not well suited for the study of dynamic phenomena, such as free and confined diffusion or active transportation of biomolecules, due to the limited scanning speed of the excitation beam across the field of view. For the same reason, confocal microscopy and similar techniques based on laser scanning are generally not used in SML microscopy because of the long acquisition times, whereas wide-field imaging configurations are usually preferred.

A good compromise between speed, contrast, and penetration depth is achieved in light sheet microscopy (LSM), a wide-field technique that employs an objective to create a thin sheet of light that selectively excites the sample plane, while a second objective collects the emitted fluorescence in the perpendicular direction [4]. The application of light sheet microscopy to SML microscopy and tracking, however, shows some important limitations. On one side, the use of two objectives strongly constrains the geometrical design of the sample chamber because of the physical dimensions of this kind of lenses. Also, specific optical arrangements are needed to avoid distortion of the excitation beam and uniform illumination of the sample in proximity of the sample-coverslip interface. A number of geometries and custom-designed sample chambers have been proposed to address these limitations [5]–[16], which are extensively reviewed elsewhere [17], [18].

A simpler approach for this kind of applications are single-objective LSM configurations, which can be easily implemented on a standard inverted fluorescence microscope to create an oblique sheet of light that illuminates cells attached on a coverslip [19]–[22]. Single-objective oblique illumination was first proposed by Tokunaga et al. in 2008 [19], with the name of HILO (Highly Inclined and Laminated Optical sheet) microscopy. In this work, a thin sheet of light was created upon refraction at large angles at the glass-water interface by focusing a Gaussian beam at the back aperture of a TIR (Total Internal Reflection) objective, laterally displaced from its optical axis (Fig.1a). With this simple approach, the authors reported a beam thickness of about 7 μm, with a 16 μm diameter Field-Of-View (FOV) and a 7.6 folds improvement in the signal-to-background ratio (SBR) compared to standard epifluorescence configuration.

**Figure 1.**
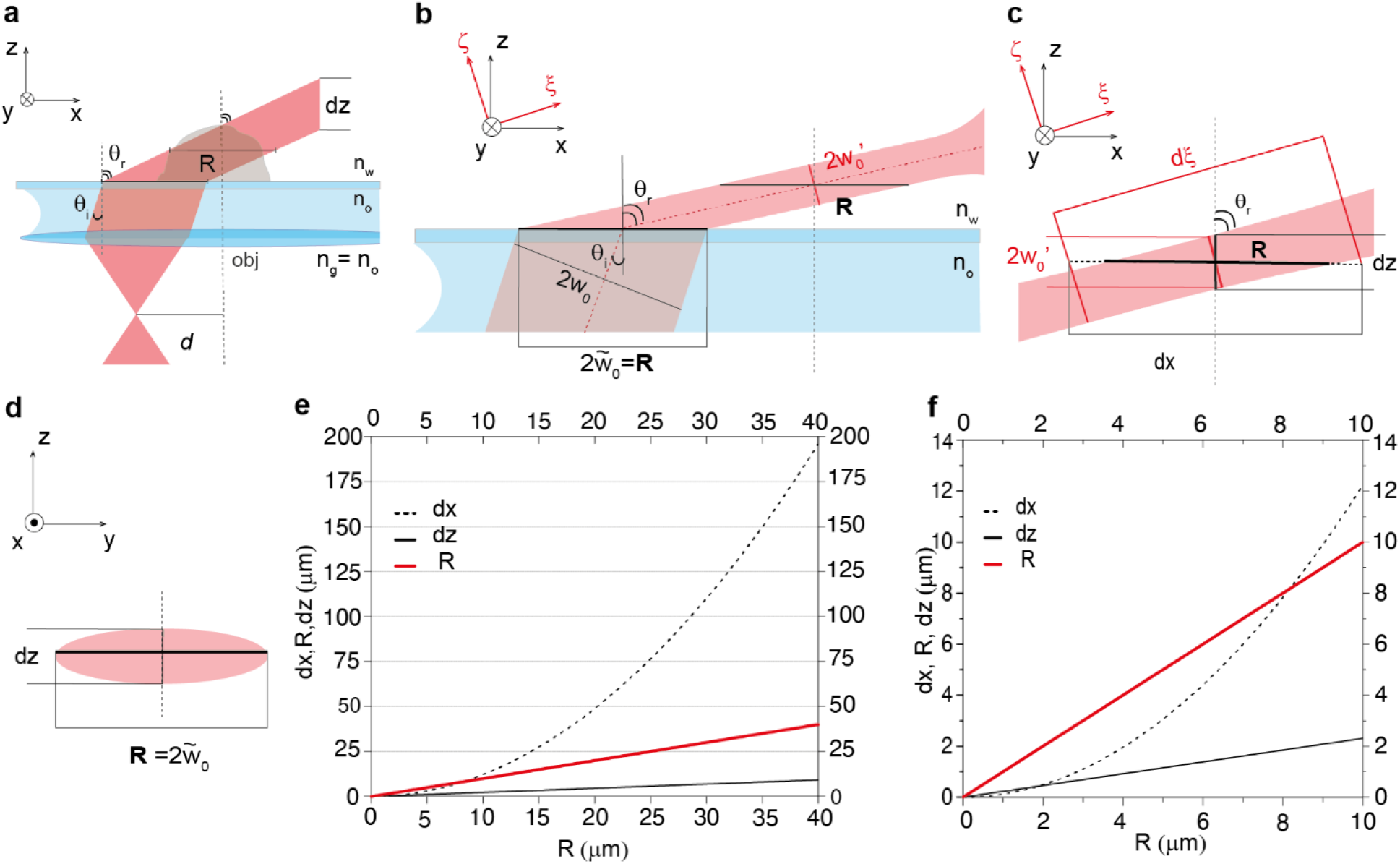
**a)** Optical scheme of highly inclined illumination according to geometrical optics. A collimated laser beam is focused in the back focal plane of a TIRF objective at a distance *d* from its optical axis (dotted grey line). The beam out of the objective is collimated and tilted by an angle *θ*_*i*_ and reaches the glass/water interface without any further deviation thanks to the optical coupling between the lens objective and the immersion oil. Upon refraction at the glass/water interface the beam is further inclined by an angle *θ*_*r*_ following Snell’s law. *n*_*g*_, *n*_*o*_, *n*_*w*_ are refractive indexes of glass, immersion oil and water respectively. *R* is the diameter of the illuminated area at the specimen plane, *θ*_*r*_ is the angle of refraction and *dz* is the thickness of the inclined beam along the objective optical axis, which are linked together by the relation *dz* = *R*/*tanθ*_*r*_. **b)** Optical scheme of the inclined beam according to Gaussian optics. Equations 1.1–1.4 are obtained by projecting a Gaussian beam propagating in the reference frame (*ξ*, *υ*, *ζ*) onto the reference system of the microscope *(x, y, z)*. **c), d)** Detailed representations of the main parameters expressed in equations 1.2–1.4 in the *(Ox,Oz)* and *(Oy,Oz)* planes respectively. **e)** Plot showing the relationship between parameters *R*, *dx*, and *dz* for *θ*_*r*_ = 77 ° from equations 1.2–1.4 for 0 < *R* < 40 μm. **f)** Plot showing the same relationship in e) restricted to 0 < *R*<*10* μm.

Despite the widespread use of inclined illumination as a simple method to reduce the fluorescence background in wide-field imaging and SML [23]–[31], proper modelling of the beam propagation, together with a simple experimental characterization of the actual inclined beam profile and performance in super-resolution microscopy are still missing.

In this work we frame a simple model based on Gaussian beam propagation theory to predict the three-dimensional shape of the inclined beam at the sample and the accessible FOV. We validate the model through both far- and near-field measurements that could be readily adopted by the users to precisely set the inclined beam size and alignment. In addition, we propose a straightforward method to reduce the beam size to sub-cellular thickness, while retaining a FOV sufficient for single cell imaging and we quantify the effects of a progressive reduction of the beam size on the quality of diffraction-limited images. Moreover, we give detailed indications to arrange an optical setup to easily control the inclined beam shape and angle according to the experimental needs. Finally, quantify the impact of the inclination of the excitation beam and of the progressive reduction of its thickness on super-resolution PALM/STORM images quality. In particular, we quantify the effects on the number of localizations and the localization precision, which are the two parameters mainly determining the resolution of the image, for two widely used, spectrally well separated, chromophores: Alexa Fluor 488 (AF488) and Alexa Fluor 647 (AF647).

## RESULTS and DISCUSSION

### Inclined Gaussian beam propagation model

Single-objective oblique illumination is implemented through a oil-immersion TIR objective by focusing the incident laser beam at the back focal plane of the objective at a distance *d* from its optical axis (Fig.1a). The beam emerging from the objective lens is collimated and propagates inside the immersion oil and glass coverslip inclined by an angle *θ*_*i*_ (we assume the oil refractive index n_o_ equal to the glass refractive index n_g_). When *θ*_*i*_ is smaller than the critical angle at which total internal reflection occurs, the beam is refracted by an angle *θ*_*r*_ at the glass-water interface and compressed along the optical axis z as a laminated sheet of light that propagates inside the sample volume (Fig.1a).

According to geometrical optics, the thickness of the refracted light sheet *dz* along the *z* direction is given by *dz* = *R*/*tanθ*_*r*_, where *R* is the diameter of the illuminated area at the specimen plane (Fig.1a). Thus, the thickness of the light sheet can be reduced by either increasing *θ*_*r*_ or decreasing *R*. However, when *R*/(*tanθ*_*r*_) is less than a few tens of microns, the beam divergence becomes appreciable and geometrical optics does not accurately describe light propagation anymore.

A better description of the beam propagation is given by Gaussian optics. We model the excitation beam as a collimated Gaussian beam that, after focusing on the back aperture of the objective lens at a distance *d* from the optical axis, impacts the glass-water interface at the front focus of the objective with a waist given by 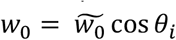 (Fig1.b). Here, 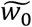 is the beam waist at *θ*_*i*_ = 0 (*d*=0). Upon refraction at the glass-water interface, the beam is tilted by and angle *θ*_*r*_ with respect to the optical axis and restricted to a beam waist 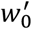 given by [32]

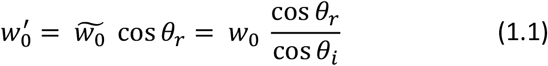

Therefore, the higher *θ*_*i*_ (and *θ*_*r*_), the thinner 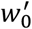.

As shown in Figure 1c, the thickness *dz* of the refracted beam along the optical axis is given by

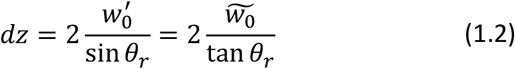

while along the *y* axis the beam is unaffected by refraction (Fig.1d), thus setting the beam size *dy* to

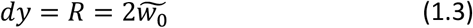

Equations 1.2 and 1.3 are formally identical to geometrical optics equations where the beam size is replaced by 2 times the beam waist. However, since the beam diverges from its waist, it is important to evaluate the distance *dx* over which the beam intensity remains approximately constant within the field of view, which is directly related to the variation of the beam thickness along the direction of propagation ξ (Fig 1b,c). The confocal parameter of the beam (*ξ*_0_) is defined as the distance over which the light sheet thickness is no greater than a factor of 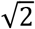 with respect to its waist, thus we can get the projection of *ξ*_0_ along the *x* axis as

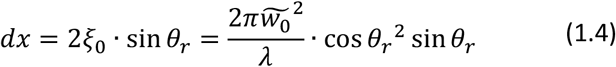

and account it as the usable width of the light sheet in the focal plane where the intensity of the inclined beam can be considered uniform.

To optimize the image quality for subcellular scale measurements, a compromise between high sectioning capability (i.e. small *dz*) and large field of view (i.e. large *dx* and *R*) should be found by adjusting the angle of refraction *θ*_*r*_ and the field of view R. On one hand, higher refraction angles are optimal to reduce the light sheet thickness, on the other hand the increased back reflected light at the glass-water interface decreases the intensity of the light refracted inside the sample and increases background fluorescence from the coverslip surface, thus deteriorating the image quality. We found that an optimal compromise is obtained by setting *θ*_*r*_ = 77°, where the intensity of the refracted beam reaches values of 88.6% and 79.9 % of the total intensity carried by the incident beam for p- and s-polarized light, respectively (see Supporting Info)[33]. Figures 1e,f show the relationship between the parameters *R*, *dx*, and *dz* for *θ*_*r*_ = 77 ° from equations 1.2, 1.3, and 1.4. For R > 8 μm, *dx* is always greater than *R*, which assures uniform illumination of the sample within the whole FOV, while, for *R* < 8 μm, *dx* is smaller than *R* and the uniformity of the illumination intensity is not guaranteed over the whole FOV anymore.

Finally, from the definition of the numerical aperture *NA* = *n*_*oil*_ sin *θ*_*i*_ and of the objective back aperture *D* = 2 *NA f*_*obj*_ [34], we can get *d* = *n*_*oil*_ *f*_*obj*_ sin *θ*_*i*_ and, thus

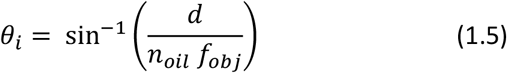

While, from Snell’s law we get

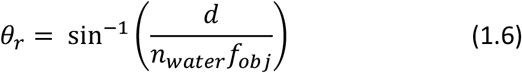

Equations 1.5, 1.6 relate *θ*_*i*_, *θ*_*r*_ to *d*, thus are of particular relevance to experimentally adjust the angle of the inclined light sheet by displacing the incoming beam from the optical axis at the objective entrance, as described in detail in the following sections.

### Measuring the light sheet inclination and dimension in the far field

The Gaussian beam model previously described can be tested experimentally by imaging a transverse section of the refracted beam in the far field, i.e. at a distance much larger than the light wavelength (643 nm in our case). Such a procedure is also a valuable way to finely adjust the beam alignment and calibrate the refraction angle *θ*_*r*_ as a function of the beam displacement *d*. For this procedure, the incident beam is focused at the back-focal plane of the objective at a distance *d* from the optical axis and it is refracted at the interface between a standard glass coverslip and air (Fig.2a).

A dark screen is positioned vertically at a distance *l* from the objective optical axis and the projection of the beam intensity profile on the screen is captured using a reflex camera (Fig. 2a and Fig. S1). According to the angle of refraction, the center of the refracted beam will sit at a certain height *h* on the screen. The widths of the beam along *y* and ζ (i.e. the directions orthogonal to the beam propagation direction ξ) at a distance 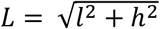 are given by

**Figure 2.**
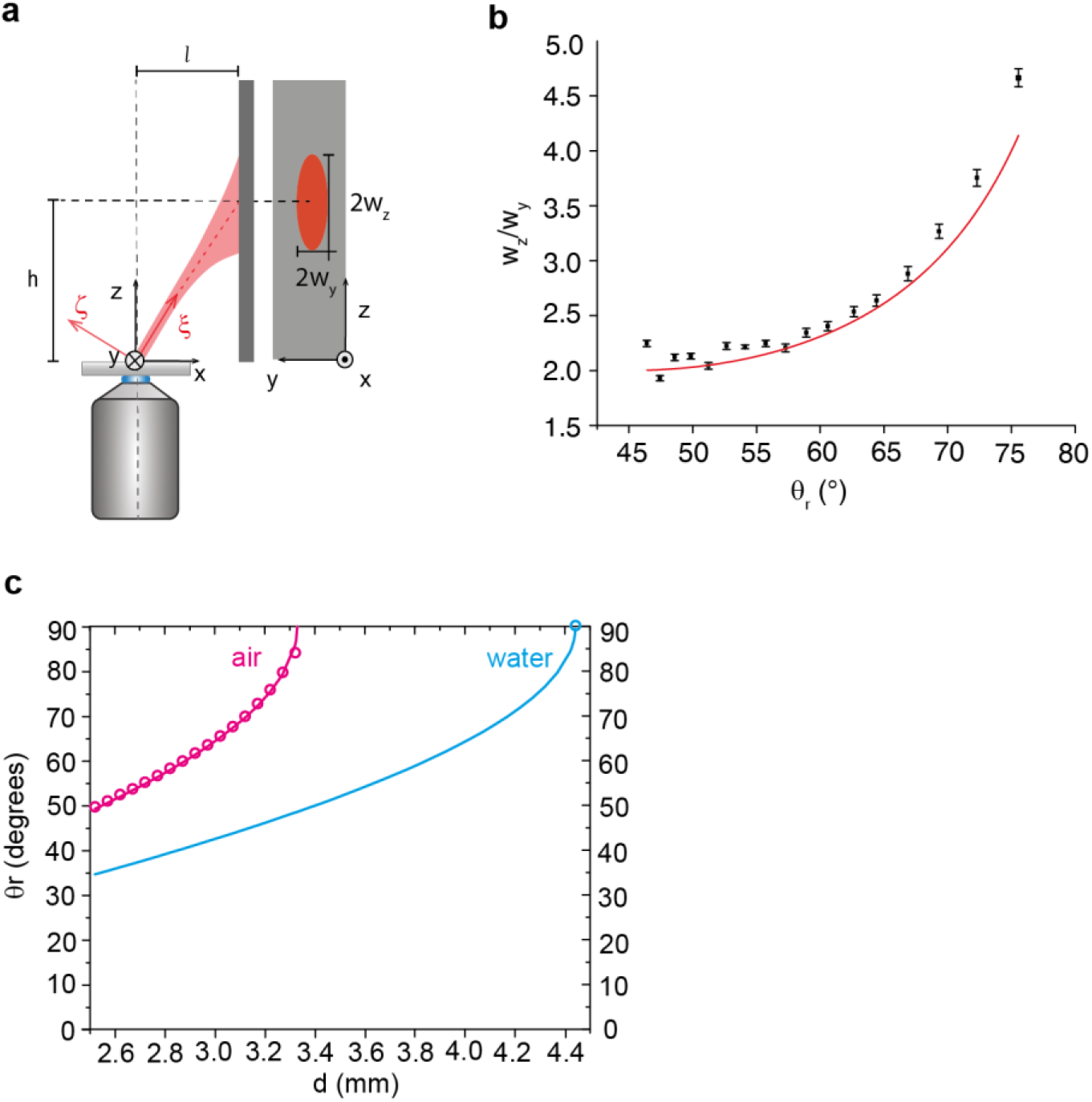
**a)** Far-field measurement. The beam inclined upon refraction at the gass/air interface is projected on a dark screen positioned at a distance *l* = 13.5 *cm* from the optical axis of the objective (see Supporting Information). The geometry of the beam is drawn in both *(Ox,Oz)* and *(Oy,Oz)* planes. *w*_*z*_ and *w*_*y*_ are the major and the minor axis respecteively of the elliptic profile projected on the screen in the *(Oy,Oz)* plane. **b)** Comparison between values of *w*_*z*_⁄*w*_*y*_ obtained from fitting the profiles on the screen with functions expressed in equations 1.10, 1.11 (black dots, error bars are errors on the fitting parameters) and values predicted from equation 1.12 (red curve). **c)** Plot of the refraction angle *θ*_*r*_ as a function of the distance *d* from the optical axis of the objective. Magenta dots are experimental values obtained from equation 1.13, while magenta and cyan curves are theoretical values predicted by equation 1.6 with *n* = *n*_*air*_ and *n* = *n*_*water*_ respectively. Blue dot is the measured distance *d* corresponding to the critical angle *θ*_*r*_ = 90° at which total internal reflection occurs.

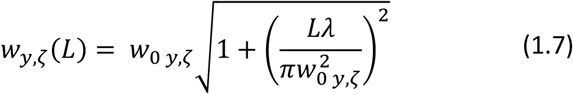

Where

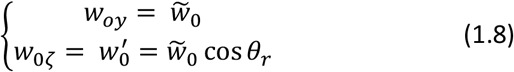

For 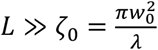, the first term in equation (1.7) can be neglected, thus giving

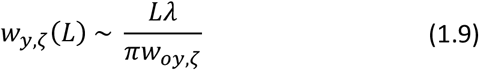

The intensity profile is projected on the screen along the y and z axis, with widths *w*_*y*_(*L*) and *w*_*z*_(*L*) = *w*_*ζ*_(*L*)/ sin *θ*_*r*_ respectively. The intensity profiles acquired on the screen can then be fitted with the following functions:

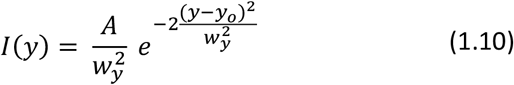

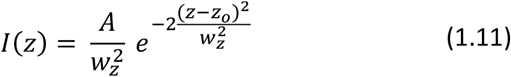

to get *w*_*z*_ and *w*_*y*_. From equations 1.8, 1.9 it comes that the ratio between *w*_*z*_ and *w*_*y*_ is solely dependent on the angle of refraction as follows:

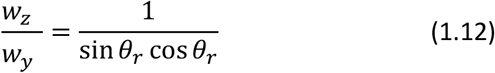

Figure 2b shows the ratio *w*_*z*_⁄*w*_*y*_ measured at different angles of refraction in air (black dots) and compared with what predicted from equation 1.12 (red curve).

From the far field images on the screen, we could measure the angle *θ*_*r*_ as

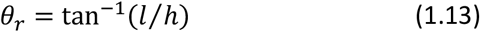

and plot its dependence on the distance *d* of the excitation beam from the optical axis of the objective. Figure 2c shows *θ*_*r*_ measured in air as a function of *d* (magenta circles) compared with values predicted from equation 1.6 with *n=n_air_* (magenta curve).

The good agreement between predictions and measurements for both *w*_*z*_⁄*w*_*y*_ (Fig.2b) and *θ*_*r*_ (Fig.2c) validates our model and enables to predict the value of *θ*_*r*_ as a function of *d* for water immersed samples, such as adherent cells, from equation 1.6 with *n = n_water_* (cyan curve in figure 2c). Moreover, we verified the distance *d* corresponding to the critical angle *θ*_*r*_ = 90°, i.e. to total internal reflection at the glass-water interface, by finding the distance from the objective optical axis at which the maximum back reflected intensity was measured (blue circle in Fig. 2c).

Finally, we set an optical system to adjust *θ*_*r*_ very precisely thanks to controlled motorized alignment of the optical components to finely change the distance *d* of the excitation beam from the optical axis to reach the desired inclination of the light sheet in the sample according to equation 1.6. The optical system is detailed in the Methods and supplementary figure S3.

The combination of ad hoc beam shaping through different Beam Stops and precise angle regulation allows to adjust the illumination condition in a highly controlled versatile and straightforward manner and could be easily implemented on any wide-filed microscope.

### Measuring the beam dimension in the near field

To experimentally verify the predicted size and shape of the inclined beam at the sample plane and optimize it under typical experimental conditions for adherent cells imaging, we set up a method for direct measurement of the inclined beam profile inside a synthetic sample with refractive index close to that of water. A 2% agarose gel with dispersed 100 nm diameter fluorescent beads (see Methods) was optically scanned through the optical arrangement shown in figure 3a. By displacing lens L5 in the detection path by means of a micrometer translation stage (Fig. 3a), different planes within the sample volume were imaged onto the EMCCD camera sensor consecutively (see Supporting Information). Fluorescent beads were excited by the light sheet inclined at *θ*_*r*_ = 77° and imaged at different depths from the glass coverslip surface up to 8 μm inside the sample volume in 1 μm steps in a “to-and-fro” modality [19] (see Supporting Information).

**Figure 3.**
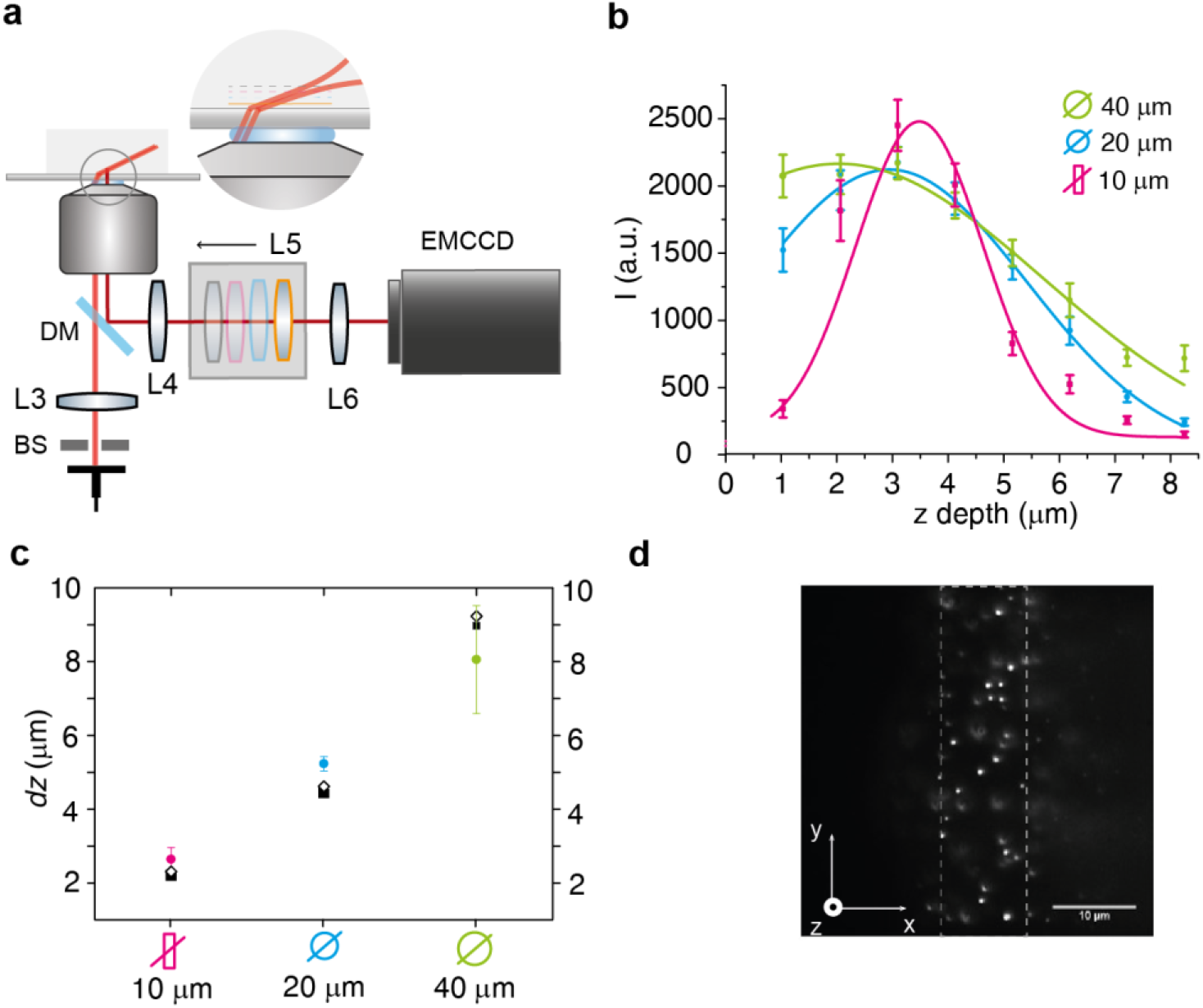
**a)** Optical scheme of the detection scanning system. The lens L5 is displaced by known distances (see Supporting Info for calibration) by means of a micrometer translator while the objective lens is fixed. At each position of L5 a different plane within the sample volume is imaged onto the EMCCD camera sensor. Images of fluorescent beads embedded in 2% agarose were taken at different depths, from the glass coverslip surface up to 8 μm inside the sample volume in 1 μm steps to reconstruct the inclined beam profile along the optical axis *z*. **b)** Average fluorescence intenisty of beads as a function of depth (*z*) with *θ*_*r*_ = 77° and *R* = 40 μm (green), *R* = 20 μm (cyan) and R[x,y]= 10 μm,40μm (magenta) respectively. The intensity at each depth is the average of *</>* (from equation S1.12 in Supporting Info) between beads belonging to the same plane. Error bars are the standard deviation of the mean. Green, cyan and magenta curves are Gaussian function fits of the experimental data. **c)** Comparison of *dz* calculated as FWHM of Gaussian fits. Coloured circles are FWHMs from fitting of profiles in b): 8 ± 1 μm (R = 40 μm, green), 5.2 ± 0.2 μm (R = 20 μm, cyan), 2.6 ± 0.3 μm (R[x,y]= 10 μm,40μm, magenta). Error bars are errors on the fitting parameters. Black squares are FWHMs from fitting of simulated profiles: 8.98 ± 0.007 μm (R = 40 μm), 4.45 ± 0.02 μm (R = 20 μm), 2.2047 ± 0.0005 μm (R[x,y]= 10 μm,40μm). Error bars are errors on the fitting parameters. Black rhombuses are values predicted from equation 1.2 (9.2 μm for R = 40 μm; 4.6 μm for R = 10 μm, 2.3 μm for R = 10 μm). **d)** Example image of 100 nm fluorescent beads excited with the beam inclined by *θ*_*r*_ = 77° and R[x,y]= 10 μm,40μm.

We then reconstructed the intensity profile of the inclined beam along the *z* axis by measuring the average fluorescence intensity of the beads over the field of view at different planes (Fig. 3b and Methods). By fitting the profile in figure 3b with a Gaussian function we evaluated the thickness of the beam *dz* as the full width at half maximum (FWHM) (Fig.3c, colored circles). The axial intensity distribution was measured for: i) *R* = 40 μm, corresponding to the full FOV in our setup (green in Fig. 3b,c), ii) *R* = 20 μm (cyan in Fig. 3b,c), and iii) R = 10 μm (magenta in figures 3b,c). *R* was adjusted through the beam stop (BS) positioned at a plane conjugated with the focal plane of the objective shown in figure 3a and supplementary figure S3a.

As reported in figures 3b,c, by reducing the diameter of the inclined beam (*R*) the thickness *dz* is reduced accordingly. Indeed, subcellular thickness (i.e. FWHM ≤ 3 μm) is achieved for *R* = 10 μm, but at the expense of the accessible FOV on the image plane, which would limit the application to single cell imaging. To mitigate this drawback for configuration iii), we shaped the beam through a linear slit BS and obtained a confined illumination with a reduced FOV along the refracted axis *x* (i.e. 10 μm), while maintaining the full FOV along the *y* axis (i.e. 40 μm). Figure 3d shows an example of fluorescent beads in agarose gel excited at *θ*_*r*_ = 77° with *R_x_* = 10 μm, *R_y_*= 40 μm (referred to as *R*[*x*, *y*] = 10 μm, 40μm from now on). Under this condition, represented in magenta in Figures 3b,c, the measured light sheet thickness was 2.7 ± 0.4 μm (FWHM of magenta profile in Fig.3b), thus suitable to enable subcellular optical sectioning.

Inn figure 3c we also compared measured versus simulated beam thicknesses *dz*. Namely, a Gaussian beam propagating at an angle *θ*_*r*_ = 77° with respect to the microscope reference frame, was simulated in Matlab based on the Gaussian optics model described above (Equations 1.1–1.4) for the three experimental conditions i), ii) and iii) described above. At each depth (8 planes separated by 1 μm starting from the glass coverslip surface through the sample), the average intensity over the FOV was calculated as the integral of the intensity over the (O*x*, O*y*) plane (see Supporting Information). By repeating the calculation for the three conditions i), ii) and iii) we obtained a set of simulated intensity profiles along the z axis analogue to the measured ones in figure 3b. The simulated profiles were then fitted with a Gaussian function to calculate the beam widths as FWHMs. Figure 3c shows the comparison between measured (colored circles) and simulated (black squares) *dz*. The good accordance between the simulated values and those from near-field measurements further validates our model and shows a good accordance also with values of *dz* predicted from equation 1.2 (black rhombuses).

### Impact of inclined illumination on diffraction limited fluorescence images quality

The effect of the inclined beam on the quality of standard diffraction limited fluorescence images was then evaluated in the three illumination conditions i), ii) and iii) on both the synthetic sample made of 100 nm fluorescent beads embedded in 2% agarose gel and on biological samples. For the evaluation on the synthetic sample, we added 2.6 nM AF647 to the agarose gel to increase the background fluorescence level (see Methods) and better evaluate the effect of the beam size reduction on the image contrast. The average Signal-to-Background Ratio (SBR) of in-focus beads was evaluated based on the fitting of the beads intensity profile through a single-molecule localization algorithm[35], as described in detail in the Methods. Figure 4a shows SBR values for conditions i), ii),iii) and for standard epifluorescence illumination (condition iv), black circle in figure 4a *x* axis label and in figure 4b and 4c legends). These measurements clearly show an improvement of the SBR with inclined illumination compared to standard epifluorescence, and clearly display a direct relationship between a reduced thickness of the inclined beam and a higher SBR in the resulting images.

**Figure 4.**
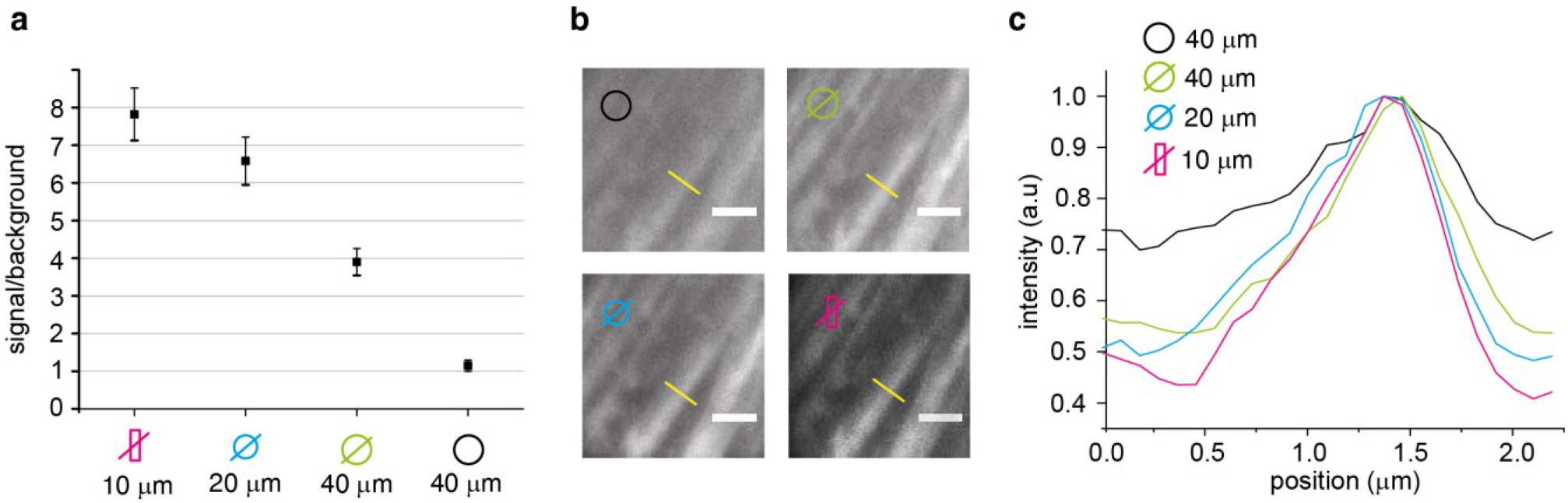
**a)** Signal-to-background-ratio (SBR) of fluorescent beads embedded in 2% agarose with dispersed 2.6 nM phalloidin-AF 647 (see Methods). Different ratios were measured in the four different illumination conditions: *θ*_*r*_ = 77°, *R* = 40 μm (green label, condition i) in the main text); *θ*_*r*_ = 77°, *R* = 20 μm (cyan label, condition ii) in the main text); R[x,y]= 10 μm,40μm (magenta label, condition iii) in the main text); wide field epifluorescence, R = 40 μm (black label, condition iv) in the main text). Data are average ± st.dev. (N = 40). **b)** Details of diffraction-limited images of the actin cytoskeleton of mammalian cells labelled with phalloidin-AF647 (see Methods) in the same four illumination conditions. Label colors as in a). The yellow line indicates the position of the transverse intensity profiles shown in c). Scale bar: 2μm. **c)** Intensity profiles of the same actin fiber, plotted across the yellow line in b) for the four illumination conditions. Label colors as in a) and b).

This trend is also confirmed qualitatively by observing images of the actin cytoskeleton of mammalian cells labelled with phalloidin-AF647 (see Methods) in the four illumination conditions (Fig.4b). The image contrast is evaluated by measuring the intensity profile across an actin stress fiber. Figure 4c shows the overlapping of the intensity profiles across the same actin stress fiber for the four illumination conditions, thus showing both an actual improvement of the contrast with inclined illumination with respect to epifluorescence and an improvement of the contrast with the reduction of the incident beam size *R*.

### Impact of inclined illumination on super-resolution images quality

Localization-based super-resolution imaging (STORM/PALM) is especially susceptible to fluorescence background, given the small amount of photons emitted by single fluorophores; as such, we evaluated the changes in the quality of super-resolution imaging in typical cell cultures. In particular, we quantified the effects of the inclined illumination in combination with beam size reduction on both the number of localizations, which affects the resolution in terms of sampling capability and determines the duration of an image acquisition, and on the localization precision, which directly affects the achievable image resolution [36]. We characterized two broadly employed photoswitchable chromophores emitting in two well separated ranges of the visible spectrum: AF488 and AF647.

The actin cytoskeleton of cultured HEK 293 cells was labelled with a phalloidin conjugate of AF488 and AF647 respectively (see Methods), and series of 20 thousand images were acquired in each of the three following configurations: i) epifluorescence *R* = 40 μm, ii) inclined light sheet with *θ*_*r*_ = 77°, *R* = 40 μm, and iii) inclined light sheet *θ*_*r*_ = 77°, *R*[*x*, *y*] = 10 μm, 40μm.

Figure 5a shows the relative number of fluorescent molecules localized by the software ThunderStorm [37] over 20 thousand frames for each configuration for both AF488 (blue squares) and AF647 (red squares). All frames were acquired on the same FOV by switching configuration every 10 thousand frames in “to-and-fro” acquisition series to account for photobleaching (10 thousand “to” plus 10 thousand “fro” frames for each configuration, see Supporting Information). The results clearly indicate that inclining the illumination beam leads to an increase in the number of localizations for both AF488 and AF647 with respect to standard epifluorescence, as a consequence of the background reduction which makes more chromophores emerge above the background threshold. Notably, at *θ*_*r*_ = 77° and *R*[*x*, *y*] = 10 μm, 40μm, the number of localizations increases 1.55 folds with respect to standard epifluorescence for AF647 and almost 2.6 fold for AF488 (Fig.5a). On the other hand, our results also point out that the inclination of the excitation beam itself, without any further size reduction, does not impact largely on the number of localizations as in the case of the beam thickness being reduced to sub-cellular dimensions (*R*[*x*, *y*] = 10 μm, 40μm and *dz* < 3 μm), especially for AF488. These relevant results demonstrate that in the case of chromophores emitting in a spectral range with high levels of fluorescence background, mainly due to cell autofluorescence, reducing the inclined excitation beam thickness to sub-cellular dimensions may improve considerably the sampling capacity and shorten the time needed to acquire PALM/STORM image stacks. This is evident from a qualitative comparison between the super-resolved images obtained by reconstructing the “to-and-fro” series of images described above (Figure 5c). Despite the number of acquired images (20K in total) is not sufficient to obtain a well sampled image of the actin cytoskeleton, the significant increase in the number of localizations for *θ*_*r*_ = 77° and *R*[*x*, *y*] = 10 μm, 40μm leads to a noticeably more detailed super-resolved image with respect to the other two optical configurations.

**Figure 5.**
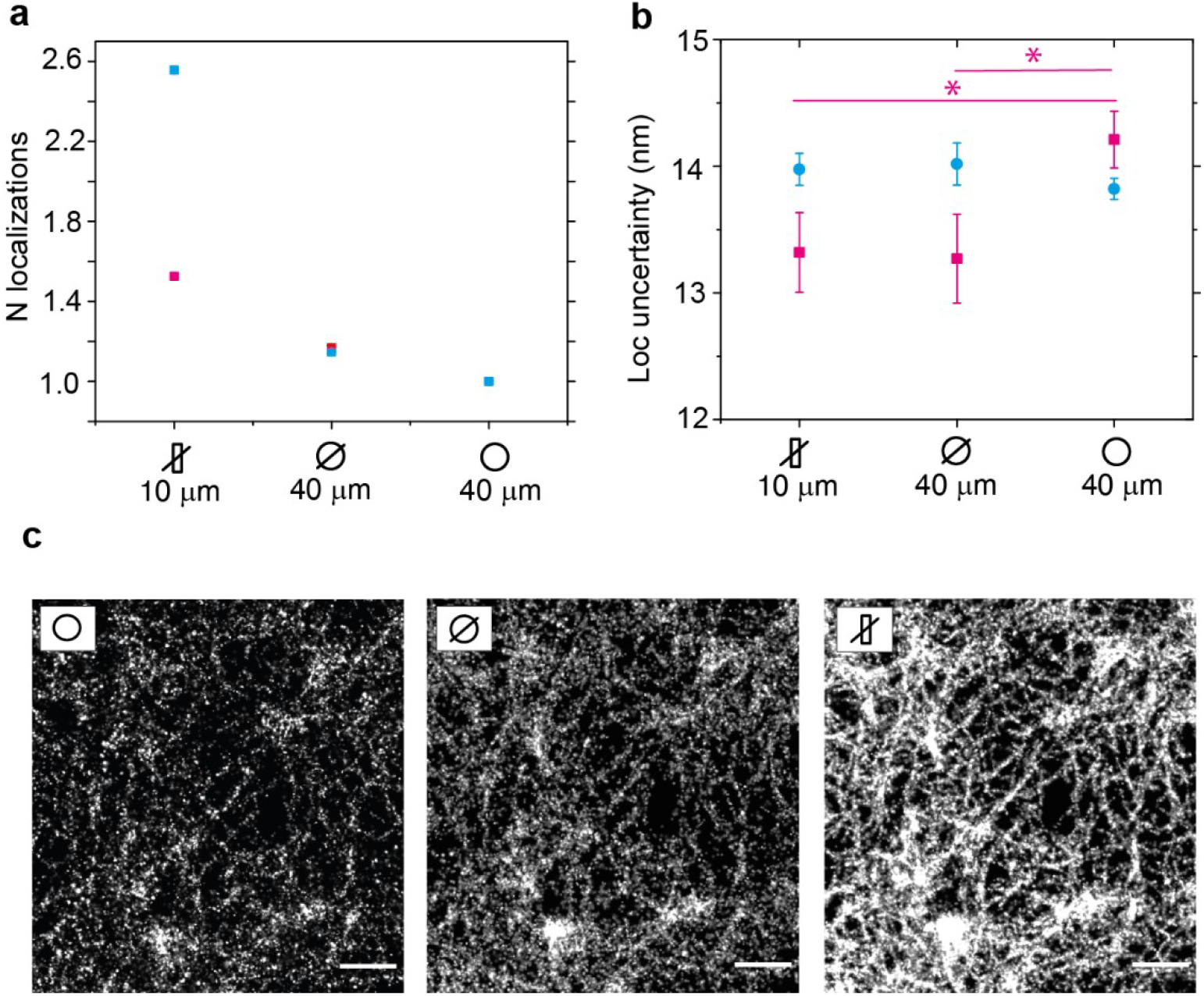
**a)** Relative number of localizations for AF488 (cyan) and AF647 (magenta) over 20 thousand frames (10 thousand “to” plus 10 tousand “fro” (see Supporting Info) in the three illumination conditions, from right to left: i) epifluorescence *R* = 40 μm, ii) θr=77°, *R* = 40 μm, and iii) θr=77°, *R*[x,y]= 10 μm,40μm respectively. The reported values are weighted averages over three indipendent measurements, normalized on epifluorescence values. **b)** Localization uncertainty for AF488 (cyan) and AF647 (magenta) in the three illuminaton configurations as in a). The uncertainty is calculated as the standard deviation of subsequent localization of persisting fluorophores (see Supporting Info). The values reported represent weighted averages between two independent experiments. Error bars are standard errors of the weighted mean. (*N*_*i*_ = 98, *N*_*ii*_ = 39, *N*_*iii*_ = 108). **c)** Super-resolved images of the actin cytoskeleton labelled with AF488 obtained from 20 thousand frames acquired in the “to-and-fro” series of images for each of the three optical configurations i), ii) and iii) respectively. The scale bar is 1 μm.

Another important parameter which is directly linked to the achievable resolution in STORM/PALM images is the localization precision. Figure 5b shows localization precision in the three optical configurations, calculated as the standard deviation of consecutive localizations of persistent chromophores (i.e. emitting for more than 8 consecutive frames) for both AF488 and AF647 (see Supporting Information). Interestingly, a significant difference is seen only in the comparison between standard epifluorescence configuration with inclined illumination conditions ii) and iii) for AF647, while no significant differences are seen for AF488.

## CONCLUSION

Highly inclined and laminated optical sheet (HILO) microscopy is a very simple and widely adopted excitation configuration capable of reducing unwanted out-of-focus signal in wide-field fluorescence imaging. HILO can reach signal-to-noise ratios close to that obtained by total internal reflection fluorescence (TIRF) microscopy. However, contrary to TIRF, HILO retains the possibility to image chromophores deep inside a sample, which made HILO a widespread technique. Despite this, a comprehensive characterization of the key parameters that determine the geometry of the inclined beam and a detailed quantification of the achievable improvements in terms of quality of both diffraction-limited and super-resolved fluorescence images was missing. In this work, we set a simple model based on Gaussian optics, which describes the key parameters of the inclined excitation beam, i.e the beam thickness *dz* and the projected confocal parameter *dx*, as a function of the beam size *R* and the angle of refraction *θ*_*r*_ (eq. 1.2, 1.3, 1.4). The use of Gaussian optics allows us to predict the beam divergence and the width of the uniformly illuminated FOV. The model was thoroughly validated by far-field and near-field measurements and simulations. The good agreement between the measured and predicted beam waist ratio *w*_*z*_⁄*w*_*y*_ (eq.1.12) and the angle of refraction *θ*_*r*_ in the far-field in air (eq.1.13) (Fig.2b,c), made it possible to predict the angle of refraction *θ*_*r*_, divergence and size of the excitation beam in a water sample in the near-field. The latter was validated from measurements of the beam thicknesses in the near-field, which were in good agreement with simulations based on our model (based on equations 1.1–1.4) (Fig.3b,c). These results make our model a “ready-to-use” pool of equations to decipher the main features of the inclined beam at the sample level, such as the thickness *dz*, and the projected confocal parameter *dx* (Fig.1e,f), which are directly linked to the quality of the acquired images and should be carefully adjusted according to experimental needs. Therefore, we propose that the far-field measurements could be adopted as a simple experimental procedure to test the correct alignment of the excitation beam, i.e. the beam angle of refraction, divergence, and size, and used to adjust the optical components until the beam in the far-field fits the model predictions. Moreover, we give indications on how to implement an optical setup to precisely set the angle of refraction *θ*_*r*_ through stepping motors, to adjust the illumination area through beam shaping and to easily switch between standard epifluorescence, inclined and eventually TIRF illumination.

Finally, we quantified the impact of the inclined beam size on both diffraction-limited and super-resolved images. In both cases we observed an improvement of the image quality with the reduction of the beam size, as a consequence of the reduction of background fluorescence from out-of-focus planes. In the case of super-resolution PALM/STORM images, we focused on the two main parameters that influence image resolution: the number of localizations, which is linked to the sampling capability, and the localization precision which sets the lower limit to the achievable image resolution. While the localization precision is only slightly improved by the reduction in the inclined beam size (Fig.5b), a significant effect was observed on the number of localized molecules per frame. In particular, we showed that, for both chromophores AF488 and AF647, the inclination of the beam leads to a significant increase in the number of localization per se (Fig.5a), while a much more substantial effect is seen with a further reduction of the beam thickness *dz* to sub-cellular dimensions, such as for *R*[*x*, *y*] = 10 μm, 40μm and *dz* < 3 μm, with an increase up to 2.6 folds with respect to standard epifluorescence and up to 2 folds with respect to inclined illumination with no further beam size reduction.

In conclusion, we provide a set of experimental and computational tools to optimize the setup and alignment of highly inclined illumination, we characterize the inclined beam geometry, signal-to-background ratio improvement, and performances in super-resolution microscopy. We also provide a simple optical arrangement to significantly increase the number of localizations allowing for better sampling capability and shorter acquisition times in PALM/STORM experiments. Finally, we share a Matlab script to easily compute the beam profile in the sample plane as a function of the experimental parameters (Supporting Information).

## METHODS

### Optical setup

The optical setup (supplementary figure S1a) is composed by an inverted wide-field fluorescence microscope (Nikon ECLIPSE TE300) equipped with 643 and 488 nm laser sources for fluorescence excitation and a 405 nm laser source for activation. The laser beams are coupled together by dichroic mirrors (DM1, DM2) and magnified 10 times by an achromatic doublet telescope L1-L2. The excitation beam is reflected by the mirror M and focused onto the back focal plane of the objective (Nikon 60X, oil immersion, NA 1.49 TIRF) by lens L3. The mirror M and the lens L3 are both mounted on a motorized translator (Fig.S3b) to displace the excitation beam at different distances *d* from the optical axis of the objective and change between epifluorescence, highly inclined, and TIR, according to equations (1.4, 1.5), see supplementary figure S1b. The size of the incident excitation beam (*R*) is regulated by a variable beam stop, BS in figure 3a and S1a (whether an iris or a linear slit) conjugated with the image plane of the objective. The fluorescence emitted by the chromophores in the sample is separated from the excitation by the dichroic mirror DM3 and projected onto the EMCCD camera (Andor iXon X3) by the tube lens L4, followed by a 3X telescope (L5, L6) necessary to obtain a pixel size suitable for single molecule localization with nm precision [2][1]. Full field of view is 40×40 μm^2^ wide, with 91 nm pixel size.

### Preparation of the calibration agarose gel with fluorescent beads

2% agarose was prepared from agarose powder (SIGMA A9539) and Milli-q ultrapure water. The solution was melted in a microwave oven and kept at about 60 °C by immersion in a thermal bath. Fluorescent TetraSpeck beads, 100 nm diameter (T7279, ThermoFisher), were diluted 10000 times in warm 2% agarose, by careful mixing. A final volume of 50 μl of this solution was sandwiched between a glass microscope slide and a glass coverslip. After the gel solidified, the fluorescent beads were embedded at different sites within the gel as shown in Fig. 3d. The bead concentration was adjusted to have them spread far apart enough to be localized through PSF fitting [1][35]. For the SBR measurements (Figure 4a) phalloidin-AF647 was diluted in the melted 2% agarose at a concentration of 2.6 nM in addition to TetraSpeck beads, to enhance the fluorescence background level.

### Beam intensity profile measurement

The profile of the inclined beam along the optical axis was measured by optically scanning the calibration sample described in the previous section while keeping the 643 nm excitation laser fixed in place. Different sample planes were focused on the camera detector by varying the position of lens L5 in the detection path through a micrometer translator (Fig. 3a, S2a, see Supporting Information).

The average intensity of in-focus beads was measured over the field of view at 8 different planes along the z axis, at 1 μm steps starting from the coverslip/sample interface. The intensity of each bead was measured as the average intensity within a Region Of Interest (ROI), shaped like a disk of width *d*_*Airy*_ = 1.22 *λ*/*NA*, minus a background level, measured as the average intensity over a ring surrounding the ROI (see Fig. S4). Considering *λ* = 680 *nm* and an image pixel size of 91 nm, the ROI diameter resulted to be 6 pixels wide, while the ring was 1 pixel wide. Intensity measurements at different planes were performed in “to-and-fro” modality to correct for changes in intensity caused by photobleaching (see Supporting Information and Fig. S7). 300 ms exposure time, 50 EM gain and 1.4 mW 643 nm laser power on the sample were set for all measurements.

The average intensity of the beads of each plane, with relative standard deviation, was plotted as a function of the depth from the coverslip. The intensity profiles, shown in Fig. 3b were then fitted with Gaussian functions to obtain FWHM as an estimate of the refracted beam thickness along the z axis.

### Signal-to-background ratio measurement of fluorescent beads

For each image, the in-focus beads were analyzed using the fitting algorithm PROOF previously developed by our group [35]. The algorithm fits a two-dimensional Gaussian function on the intensity profile of individual beads and calculates the intensity (*I*) of the central peak, as well as the background level (*I*_*back*_), corresponding to the Gaussian offset. The signal-to-background ratio (*SBR*) is then calculated as:

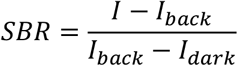

Where *I*_*dark*_ is the dark noise which was measured as the average intensity level detected by the camera in the absence of light sources for appropriate exposure time and EM gain conditions. The *SBR* was calculated for each bead, and the final *SBR* for a given illumination configuration was calculated as the average over at least 40 beads.

### Cell fixation and staining

#### STORM imaging of the actin cytoskeleton

Human Embryonic Kidney cells (HEK 293) were plated in DMEM-F12 medium with added serum and antibiotics at 10% confluency on 18 mm glass coverslips (previously coated with poly-l-lysine) and let adhere for 16-24 hours. Afterwards, cells were gently washed 3 times in phosphate-buffered saline (PBS) supplemented with 0.5 mM MgCl_2_ and 0.8 mM CaCl_2_ and then incubated in 4% paraformaldehyde (PFA) in PBS for 10 minutes. Cells were then washed with PBS with added MgCl_2_ and CaCl_2_ for 3 times and permeabilized by incubating with a 0.075% Triton X-100 solution in PBS for 7 minutes.

After 3 washes in PBS supplemented with MgCl_2_ and CaCl_2_, cells were blocked by incubating for 30 minutes in a 4% BSA solution in PBS with added MgCl_2_ and CaCl_2_. The blocking solution was then removed and cells were covered with approximately 400 μl of either 0.5 μM phalloidin-AF488 or 0.5 μM phalloidin-AF647 (Thermo Fisher Scientific) in PBS, and incubated overnight in the dark at 4°C. After overnight staining, cells were washed once with PBS before being mounted on an imaging chamber and immersed in 500 μl of Imaging Buffer (IB: 100 mM β-MercaptoEthylamine hydrochloride (MEA, Sigma-Aldrich), 3% (v/v) OxyFlour™ (Sigma-Aldrich), 20% (v/v) of sodium DL-lactate solution (Sigma-Aldrich), in PBS, pH adjusted to 8–8.5 with NaOH). The Imaging Buffer we employed is a slight variation on the OxEA buffer developed by Nahidiazar *et al*.[38] and is suitable for STORM imaging with both AF488 and AF647. Finally, the chamber was mounted on the microscope immediately after adding the Imaging Buffer.

#### Diffraction limited images of the actin cytoskeleton for contrast evaluation

AC26 human cardiomyocites were grown and stained with 0.5 μM phalloidin-AF647 (Thermo Fisher Scientific) by following the same procedure described above for STORM imaging.

### Image acquisition

#### Diffraction-limited images

the sample was excited with 643 nm laser beam at about 10 W/cm^2^ on the sample, with 40 ms integration time and no EM gain.

#### Direct STORM images

excitation laser (either 643 nm or 488 nm) was shined at > 1 kW/cm^2^ intensity on the sample, with 40 ms integration time and 400 EM gain. For evaluation of the number of localizations and the localization precision, continuous excitation with the 488/643 nm laser in the epi configuration was carried on for 600 seconds prior to acquisition, at high intensity (~5.3 kW/cm^2^), to reach a photoswitching steady state[36].

## Supporting Information

The Supporting Information is available free of charge at://https:

## Authors Information

M.C. and L.G. conceived the project and developed the Gaussian optics model. L.G., M.C. built the setup. L.G., V.C., M.C. performed far-field measurements. T.V., V.C. performed and analyzed near-field experiments. V.C. wrote the MATLAB code for the inclined beam intensity simulation. T.V. performed super-resolution experiments and wrote MATLAB routines for performance analysis. L.G. wrote the paper with input from all authors. M.C. and F.S.P. provided general supervision of the project.

## Acknowledgements

This work was supported by the European Union’s Horizon2020 research and innovation program under grant agreement no 871124 Laserlab-Europe.

## Data availability

Data supporting the findings of this manuscript are available from the corresponding author upon reasonable request.

